# RNA Polymerase II is a Polar Roadblock to a Progressing DNA Fork

**DOI:** 10.1101/2024.10.11.617674

**Authors:** Taryn M. Kay, James T. Inman, Lucyna Lubkowska, Tung T. Le, Jin Qian, Porter M. Hall, Dong Wang, Mikhail Kashlev, Michelle D. Wang

## Abstract

DNA replication and transcription occur simultaneously on the same DNA template, leading to inevitable conflicts between the replisome and RNA polymerase. These conflicts can stall the replication fork and threaten genome stability. Although numerous studies show that head-on conflicts are more detrimental and more prone to promoting R-loop formation than co-directional conflicts, the fundamental cause for the RNA polymerase roadblock polarity remains unclear, and the structure of these R-loops is speculative. In this work, we use a simple model system to address this complex question by examining the Pol II roadblock to a DNA fork advanced via mechanical unzipping to mimic the replisome progression. We found that the Pol II binds more stably to resist removal in the head-on configuration, even with minimal transcript size, demonstrating that the Pol II roadblock has an inherent polarity. However, an elongating Pol II with a long RNA transcript becomes an even more potent and persistent roadblock while retaining the polarity, and the formation of an RNA-DNA hybrid mediates this enhancement. Surprisingly, we discovered that when a Pol II collides with the DNA fork head-on and becomes backtracked, an RNA-DNA hybrid can form on the lagging strand in front of Pol II, creating a topological lock that traps Pol II at the fork. TFIIS facilitates RNA-DNA hybrid removal by severing the connection of Pol II with the hybrid. We further demonstrate that this RNA-DNA hybrid can prime lagging strand replication by T7 DNA polymerase while Pol II is still bound to DNA. Our findings capture basal properties of the interactions of Pol II with a DNA fork, revealing significant implications for transcription-replication conflicts.

---

Efficient and accurate DNA duplication is essential for the preservation and transmission of the genetic information of all living organisms, from bacteria to humans. However, DNA replication shares the same DNA substrate with transcription, another essential cellular function that occurs throughout the cell cycle. Thus, during DNA replication in the S phase of the cell cycle, a replisome may encounter a transcribing RNA polymerase (RNAP)^1–9^, resulting in a conflict that could lead to significant cellular consequences.

A replisome may encounter an RNAP moving either co-directionally or head-on, and the two orientations are known to lead to different outcomes concerning genome stability and integrity^10–12^. While both orientations are disruptive to replication, head-on conflicts are much more detrimental than co-directional conflicts ^3,9–11,13–19^. When a replisome encounters an RNAP head-on, replication progression is greatly impeded, and the replisome can be severely stalled. This may lead to replisome disassembly, fork reversal, and fork restart, creating a host of downstream effects that compromise cellular function^8,9,11,20,21^. In contrast, while co-directional conflicts are still disruptive to replication, they do not stall the replisome to the same extent as head-on conflicts^1,11,22^.

Despite the cellular consequences of these conflicts, our understanding of their underlying mechanisms remains limited. Although RNAP has been found to be a more severe roadblock to replication when approaching a replisome head-on versus co-directionally^3,9–11,13–19^, the fundamental cause for the RNAP roadblock polarity remains unclear. Furthermore, head-on conflicts, not co-directional ones, have been found to promote the formation and accumulation of R-loops, three-stranded RNA-DNA hybrid structures with the nascent RNA transcript reannealed to the template DNA strand at the region of the encounter^23–25^. However, the structure of these R-loops remains largely speculative. The prevailing view generally places the R-loop behind the RNAP in the context of head-on replication-transcription conflicts, but this view has not been validated experimentally^9,25–27^. Understanding the consequences of head-on conflicts requires a method that can elucidate the nature of these R-loops, which has proven experimentally challenging.

In this work, we have approached this problem using a simplified model system involving a mechanically progressing DNA fork and an elongating Pol II. The simplicity of this approach makes it possible for us to directly investigate the Pol II roadblock polarity and the structure of RNA-DNA hybrid. Using this approach, our findings provide significant insights into the nature of the Pol II roadblock that are relevant to understanding transcription-replication conflicts.

## Pol II is an inherent polar roadblock to a progressing DNA fork

While head-on transcription-replication conflicts are more detrimental than co-directional ones^3,9–11,13–19^, it is unclear whether RNAP is inherently a polar roadblock to replication or whether head-on conflicts are more prone to R-loop formation, which then converts RNAP into a stronger roadblock to replication. To examine the RNAP roadblock polarity, we mimicked the replisome progression by mechanically unzipping DNA through a bound Pol II using a dual optical trap equipped with a multi-channel flow cell (Fig. 1a; Supplementary Fig. 1). Previously, we demonstrated DNA unzipping as a powerful approach for mapping protein-DNA interactions^28–32^. Here, we used the DNA unzipping mapper to investigate the resistance experienced by the DNA fork when Pol II transcribes away from the fork (co-directional configuration) or towards the fork (head-on configuration).

**Figure 1.**
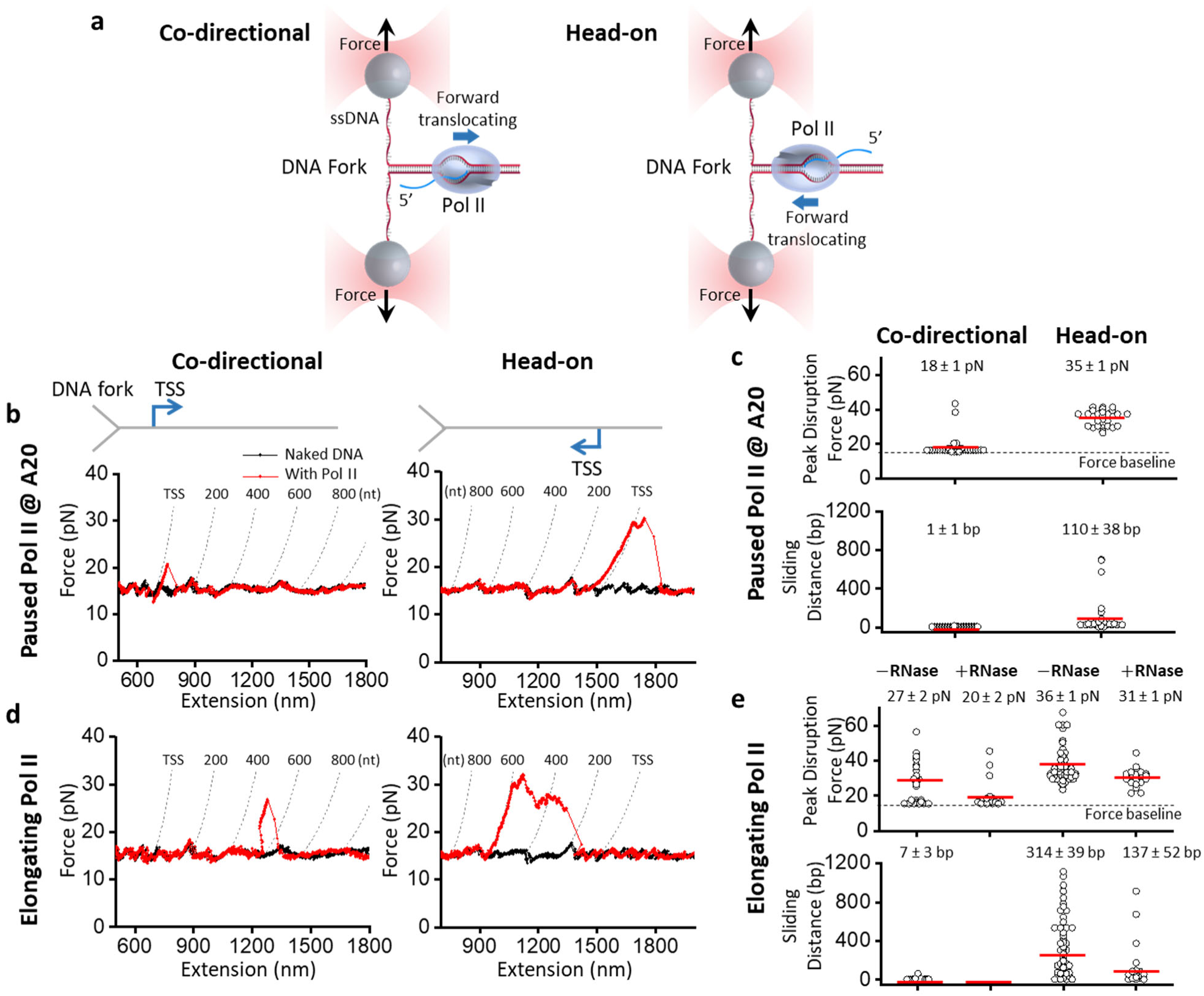
Pol II is a polar roadblock to a progressing DNA fork. **a.** Experimental configurations used to study the interaction of a DNA fork with the Pol II molecule in two collision configurations: co-directional (CD) and head-on (HO). The two daughter DNA strands are tethered between two optically trapped beads of a dual trap. The DNA fork is mechanically unzipped through a Pol II elongation complex (EC) and disrupts it. The resulting force and extension map the strength and location of Pol II interaction with DNA. **b.** Representative force-extension traces of the DNA fork unzipping through a paused Pol II EC in both collision configurations. Pol II was paused after 20 nt of RNA transcription from the transcription start site (TSS). Each dashed curve indicates the predicted force-extension curve for an unzipping fork encounter with a Pol II after transcription of the specified number of nucleotides. **c.** Scatter plots of the peak disruption force and Pol II sliding distance for both collision configurations of a paused Pol II. The mean values (also as the red bars) and their SEMs are indicated (*N* = 38 for CD; *N* = 28 for HO). **d.** Representative force-extension traces of the DNA fork unzipping through an elongating Pol II in both collision configurations. Each dashed curve indicates the predicted force-extension curve for an unzipping fork encounter with a Pol II after transcription of the specified number of nucleotides. **e.** Scatter plots of the peak disruption force and Pol II sliding distance for both collision configurations for elongating Pol II in the presence and absence of RNase T1. The mean values (also as the red bars) and their SEMs are indicated. For the peak disruption force data: CD (-) RNase, *N* = 27; CD (+) RNase, *N* = 19; HO (-) RNase, *N* = 60; HO (+) RNase, *N* = 21. For the sliding distance data: CD (-) RNase, *N* = 23; CD (+) RNase, *N* = 0; HO (-) RNase, *N* = 60; HO (+) RNase, *N* = 21.

In this experiment, we formed a Pol II elongation complex (EC) on a dsDNA template^33^, which was then ligated to two dsDNA unzipping adaptor arms (Supplementary Fig. 2)^34^. This Y-template was then unzipped with the unzipping fork approaching the Pol II either co-directionally or head-on (Fig. 1a). Using this method, we first examined a Pol II EC paused at A20 after 20 nt RNA transcription (Methods), which should have a limited length of RNA outside Pol II for R-loop formation (Fig. 1b). We found that before the unzipping fork encountered Pol II, the unzipping force followed the naked DNA force baseline; when the fork encountered Pol II, the unzipping force deviated from the force baseline. In the co-directional configuration, a bound Pol II had a mean peak disruption force of 18 pN, 3 pN above the naked DNA unzipping force baseline (15 pN), with minimal sliding along DNA under the influence of the unzipping fork (Fig. 1c). In contrast, in the head-on configuration, the unzipping force rose significantly above the baseline at the bound Pol II, with a mean peak disruption force of 35 pN, 20 pN above the baseline, indicating Pol II can significantly resist the DNA fork progression (Fig. 1c). Furthermore, the force rise persisted for about 100 bp, consistent with Pol II sliding along the DNA, resisting removal from the DNA (Fig. 1c).

These data with a Pol II paused at A20, where the possibility of R-loop formation is minimal, suggest that Pol II is inherently a stronger roadblock to a DNA fork when oriented head-on versus co-directionally. The polarity is evidenced by a significantly greater disruption force and longer sliding distance before removal. The long-distance sliding behavior was surprising given the short RNA available, suggesting that Pol II can hold on to the DNA fork even without interaction with an RNA-DNA hybrid.

To investigate whether an elongating Pol II is also a polar roadblock to the DNA fork, we allowed Pol II to transcribe on the DNA before unzipping through it (Fig. 1d). Under this condition, Pol II could transcribe a few hundred nucleotides so an R-loop could potentially form, which we systematically examine later. Here, we examined Pol II’s resistance to the DNA fork. When the DNA fork encountered an elongating Pol II moving co-directionally, the mean disruption force was 27 pN (or 12 pN above the baseline) with minimal sliding, suggesting an elongating Pol II is more stable than the paused Pol II at A20 in the co-directional configuration (Fig. 1e). When the DNA fork encountered an elongating Pol II head-on, the mean disruption force was 36 pN, 21 pN above the baseline, again greater than that of the co-directional configuration, with force rise persisting over about 300 bp (Fig. 1e). These data involving an elongating Pol II reinforce the findings from the paused Pol II and demonstrate that an elongating Pol II is also a more potent and persistent roadblock in the head-on configuration than in the co-directional configuration.

In addition, we found that an elongating Pol II is a stronger roadblock than a paused Pol II at A20 in either the co-directional or head-on configuration (compare Fig. 1c and Fig. 1e). However, it is unclear whether this roadblock enhancement results from an elongating Pol II being more stable than a paused Pol II or from a long nascent RNA that somehow allows Pol II to anchor to DNA more stably. To differentiate between the two possibilities, we performed additional experiments of an elongating Pol II in the presence of RNase T1, which can digest the RNA transcript outside the enzyme^35–37^. We found that the presence of RNase T1 significantly decreased the disruption force for both co-directional and head-on configurations. The presence of RNase T1 also considerably reduced the sliding distance, even for the head-on configuration, making such a Pol II EC behave more like a paused Pol II at A20. Thus, the presence of a long nascent RNA contributed to the enhanced stability of an elongating Pol II.

Since the progressing DNA fork used here somewhat resembles a progressing replication fork, our observations might provide a mechanistic explanation for the previous findings that head-on transcription-replication conflicts are more detrimental than co-directional ones. When a replisome encounters Pol II moving head-on instead of co-directionally, the replisome may have more difficulty removing Pol II as Pol II binds more stably to DNA and resists removal via sliding. This Pol II roadblock polarity does not require long RNA, suggesting that the polarity is inherent to a Pol II EC. Based on the Pol II EC crystal structure^38–40^ and supported by these data, Pol II interacts weakly with dsDNA behind its active site but tightly with dsDNA in front of its active site. These structural features provide a possible explanation for the Pol II polarity detected by the DNA fork and might also hold for a replisome. When a replisome approaches the Pol II from behind, the replisome first encounters the transcription bubble, which can be readily disrupted, and the disruption of the transcription bubble may destabilize the elongation complex to allow Pol II removal. In contrast, when the replisome approaches Pol II from the front, the fork first encounters the front edge of Pol II, which tightly clamps onto the dsDNA, likely making Pol II more resistant to removal.

## RNA-DNA hybrid formation

The data in Fig. 1 show that a Pol II EC becomes more stable against removal when associated with a long nascent RNA, raising the possibility that such stabilization is mediated by RNA-DNA hybrid formation. To investigate this possibility, we re-examined the co-directional unzipping traces (Fig. 1d). As shown in the more detailed view of an example trace (Fig. 2a), immediately after the unzipping fork encountered the bound Pol II and disrupted it, we detected an extension shift, where the force profile follows that of the naked DNA but at a shorter extension. In many traces, this extension shift occurs immediately upon the fork encountering Pol II (Supplementary Fig. 3). If this extension shortening was due to RNA-DNA hybrid formation, the shortening may increase with the RNA transcript length (Fig. 2b). If the entire RNA transcript forms a hybrid, the extension shortening is proportional to the RNA transcript length. Interestingly, the measured extension shortening agrees well with this prediction (Fig. 2c), indicating that the entire RNA transcript can readily form an RNA-DNA hybrid once the ssDNA becomes available near the bound Pol II.

**Figure 2.**
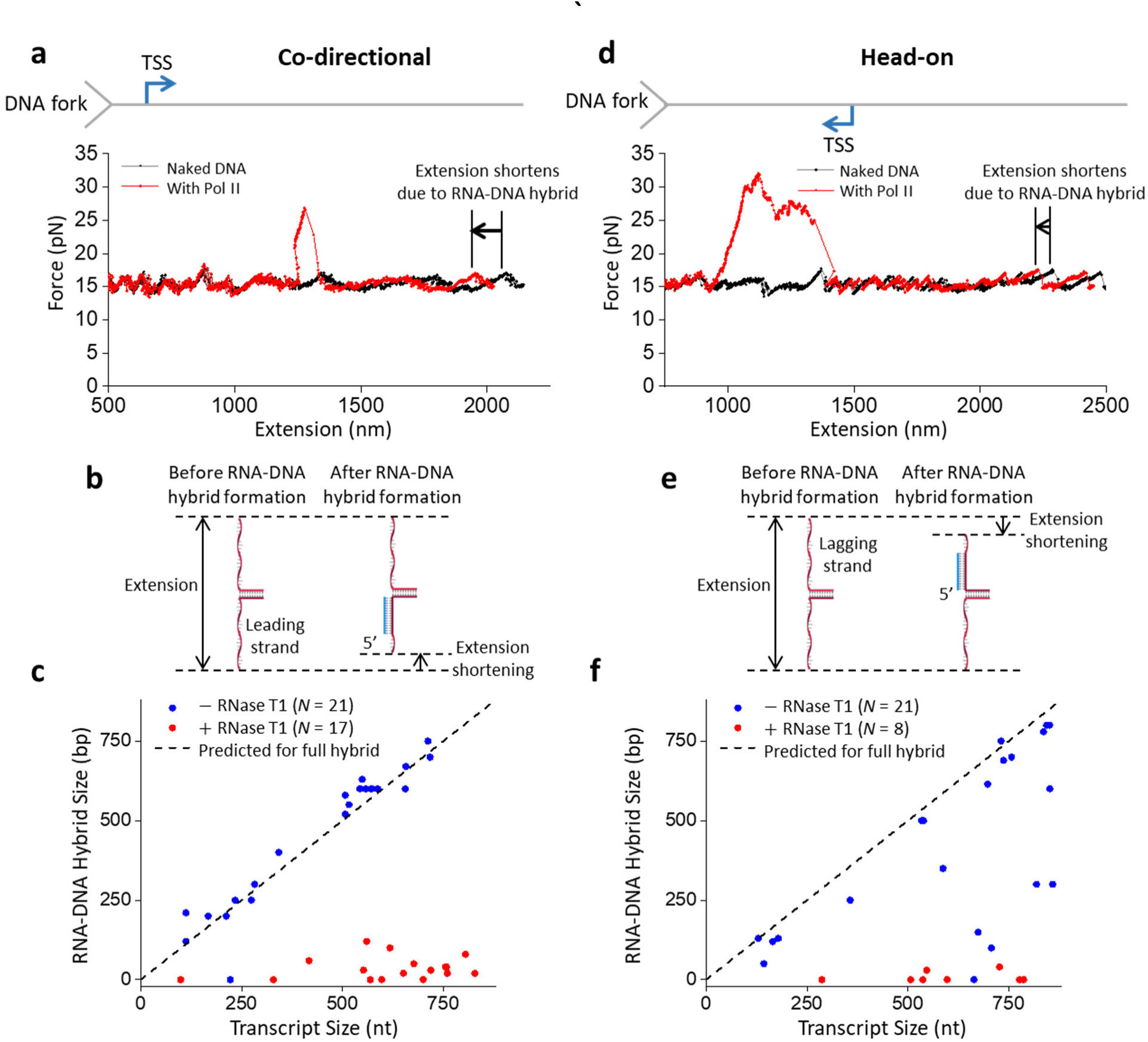
RNA-DNA hybrid formation after DNA fork collision. **a.** Detailed view of a representative force-extension trace of the DNA fork unzipping through an elongating Pol II in the co-directional configuration showing an extension shift, where the force profile follows that of the naked DNA but at a shorter extension. **b.** Cartoon depiction of how an RNA-DNA hybrid leads to extension shortening in the co-directional configuration, with the hybrid forming on the leading strand. **c.** RNA-DNA hybrid size, measured after the disruption of Pol II by the DNA fork, plotted against transcript size for the co-directional configuration. The dashed line indicates the predicted hybrid size if the entire transcript RNA is converted to hybrid. (-) RNase T1, *N* = 21; (+) RNase T1, *N* = 17. **d.** Detailed view of a force-extension trace of the DNA fork unzipping through an elongating Pol II in the head-on configuration showing an extension shift, where the force profile follows that of the naked DNA but at a shorter extension. **e.** Cartoon depiction of how the RNA-DNA hybrid leads to extension shortening in the head-on configuration, with the hybrid forming on the lagging strand. **f.** RNA-DNA hybrid size, measured after the disruption of Pol II by the DNA fork, plotted against transcript size. The dashed line indicates the predicted hybrid size if the entire transcript RNA is converted to hybrid. (-) RNase T1, *N* = 21; (+) RNase T1, *N* = 8.

We also re-examined head-on unzipping traces of an elongating Pol II such as the one shown in Fig. 1d in a more detailed view (Fig. 2d). In contrast to the co-directional encounter, the nascent RNA is located distal to the DNA fork. When the unzipping fork encountered the front of the bound Pol II (Fig. 2d), the force rose dramatically. After the unzipping force returned to the baseline, we again detected an extension shortening, where the force profile followed that of the naked DNA at a shorter extension. This shortened extension may be indicative of the extent of the RNA-DNA hybrid on the lagging strand (Fig. 2e). We found that most of the traces show a hybrid formation consistent with an entire RNA transcript being converted to the hybrid (Fig. 2f). A minority of traces had a hybrid length smaller than expected, indicating that not all nucleotides of the RNA transcript are converted to a hybrid. This suggests that some regions of the RNA may be unavailable for hybridization in this configuration.

Since RNA-DNA hybrid formation requires RNA, RNA cleavage by RNase T1 should minimize hybrid formation. Consistent with this prediction, when RNase T1 was present during transcription, DNA extension shortening after unzipping through Pol II was minimal in all traces of both the co-directional and head-on configurations (Fig. 2c,f; Supplementary Fig. 4). These observations further support that the observed DNA shortening was due to RNA-DNA hybrid formation, which can only occur in the presence of RNA.

Importantly, we never detected any RNA-DNA hybrid before the DNA fork encountered Pol II, highlighting that the hybrid formation requires the presence of ssDNA complementary near Pol II. *In vivo*, when a replisome approaches a Pol II co-directionally, ssDNA immediately behind Pol II should rarely occur until the replisome encounters Pol II. Previous studies showed that Pol II generated (-) supercoiling behind could facilitate DNA melting and R-loop formation behind Pol II^41–44^. However, such an R-loop is less likely to form and sustain when a replisome trails behind Pol II because the (+) supercoiling generated by the replisome can neutralize the (-) supercoiling from Pol II. However, if the replisome stalls at a lesion, leading to the decoupling of the replicative DNA polymerase and helicase, the continued unwinding by the helicase may generate ssDNA right behind the Pol II to enable RNA-DNA hybrid formation. Therefore, an RNA-DNA hybrid in a co-directional conflict is still possible, albeit less likely. In contrast, when a replisome encounters Pol II head-on, the replisome progression may remove Pol II but leave behind the RNA. Continued replication progression then creates ssDNA on the lagging strand for hybridization with the RNA.

## RNA-DNA hybrid in front of Pol II

Our finding that an RNA-DNA hybrid can form when ssDNA complementary is present in the Pol II vicinity raises the possibility that a hybrid may also form in front of Pol II during a head-on collision of a transcribing Pol II with a replisome *in vivo*. The replisome may be sufficiently strong to backtrack Pol II, leading to 3’ RNA protrusion from Pol II’s secondary channel^45–48^. This protruded RNA may hybridize with the lagging strand at the replication fork, where short stretches of ssDNA should be transiently present^49–51^. Thus, an RNA-DNA hybrid could form before Pol II removal in a head-on transcription-replication collision.

To investigate this possibility, we again mimicked the head-on transcription-replication collision using the DNA fork and an elongating Pol II but introduced a step to backtrack Pol II (Fig. 3a). Here, we allowed Pol II to transcribe for some distance before unzipping to Pol II. Once the DNA fork encountered Pol II, we held the unzipping force at 22 pN for 10 s to facilitate Pol II backtracking. The extent of the backtracking at this step varied from molecule to molecule, with some showing minimal backtracking (Fig. 3b) while others showed extensive backtracking (Fig. 3c). Subsequently, we attempted to rezip the DNA by reducing the extension. We observed that when Pol II showed minimal backtracking, the DNA could be fully rezipped (Fig. 3b), but when Pol II showed extensive backtracking, the DNA typically could not be fully rezipped (Fig. 3c).

**Figure 3.**
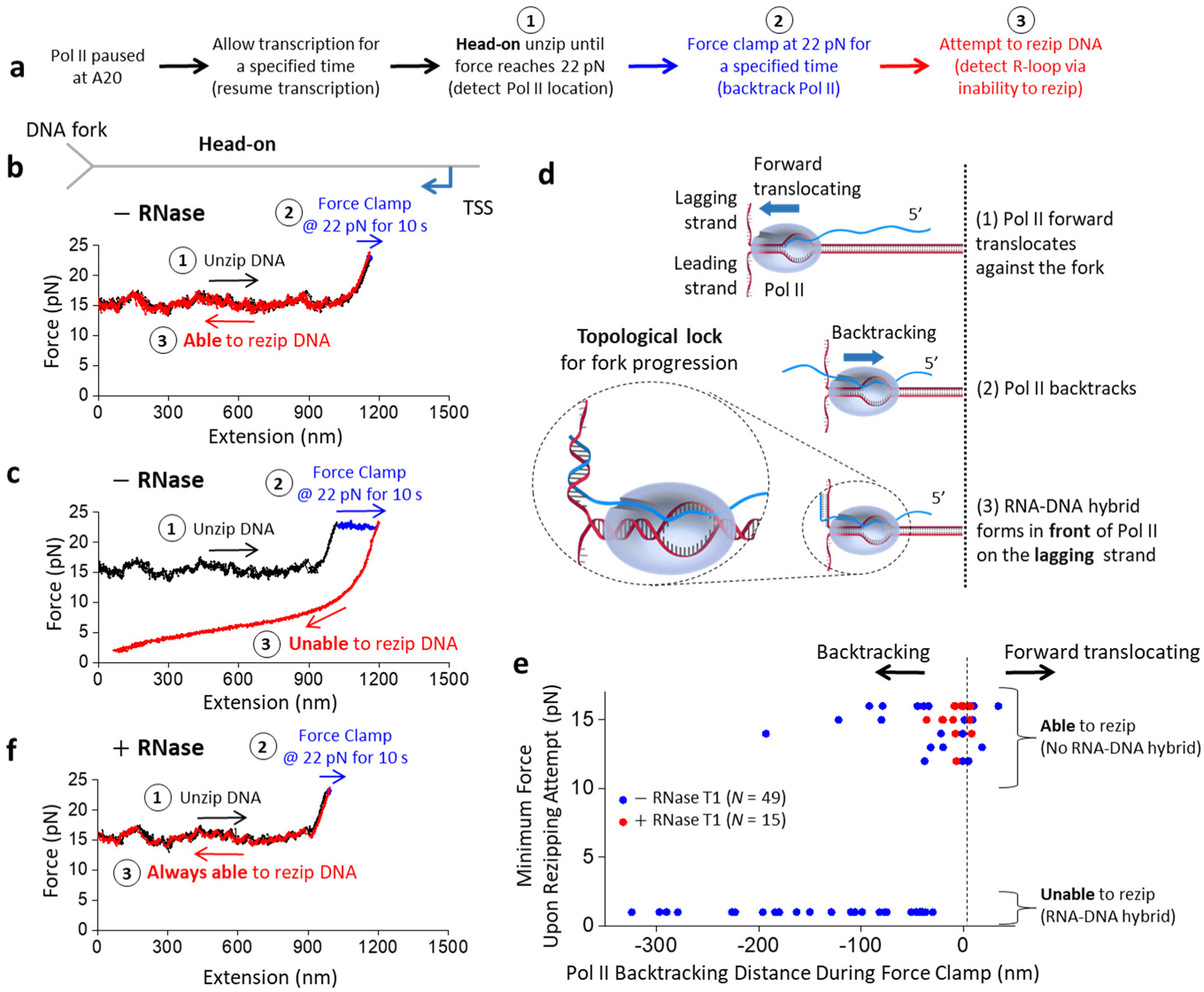
RNA-DNA hybrid formation in front of Pol II. **a.** Outline of experimental steps to backtrack Pol II and determine if RNA-DNA hybrid formation occurred. **b.** Representative force-extension trace of the DNA fork interaction with an elongating Pol II in the head-on configuration, with minimal backtracking during the force clamp step. The DNA was rezipped at step 3. **c.** Representative force-extension trace of the DNA fork interaction with an elongating Pol II in the head-on configuration, with extensive backtracking during the force clamp step. The DNA could not be rezipped at step 3. **d.** Cartoon depicting the proposed fork and Pol II conformation after Pol II is backtracked upon head-on collision with the fork. The 3’ RNA extruded from a backtracked Pol II can hybridize with the lagging strand in front of Pol II. For the ease of strand tracking, dsDNA or RNA-DNA hybrid are shown as two parallel strands, except for the inset where their helical structures are correctly depicted. The inset shows that the RNA-DNA hybrid formation restricts Pol II rotation and DNA unwinding, and topologically locks Pol II on the DNA. **e.** The minimum force reached during the attempt to rezip step versus the amount of backtracking during the force clamp step, in the presence and absence of RNase T1. A low minimum force characterizes an inability to rezip. The black dashed line indicates the initial Pol II position before the force clamp. The backtracked distance in base pairs may be estimated using a conversion factor of 1.0-1.2 bp/nm. (-) RNase T1, *N* = 49; (+) RNase, T1 *N* = 15. **f.** Representative force-extension trace of the DNA fork interaction with an elongating Pol II in the head-on configuration in the presence of RNase T1. The DNA was rezipped at step 3.

Since DNA rezipping requires both ssDNA strands to be available for base pairing, the inability to rezip indicates an obstruction in the ssDNA strands. This could occur if an RNA-DNA hybrid forms in the lagging strand in front of a backtracked Pol II (Fig. 3d). If so, then an RNA-DNA hybrid may form more readily for more extensively backtracked Pol II since there is a greater opportunity for RNA-DNA hybridization. Consistent with this expectation, we found a strong correlation between the inability to rezip (characterized by a low minimum force upon the rezipping attempt) and the backtracking distance (Fig. 3e), suggesting that the inability to rezip is an indicator of an RNA-DNA hybrid in front of Pol II.

To validate this interpretation further, we carried out the same experiments but in the presence of RNase T1, which can degrade RNA to limit Pol II backtracking (Fig. 3f). Without backtracking, the 3’ RNA will not protrude from Pol II’s secondary channel to allow RNA-DNA hybrid formation in front of Pol II. Thus, the presence of RNase should enable more efficient rezipping. Indeed, in the presence of RNase T1, we detected minimal backtracking, with the inability to rezip abolished entirely and all traces being rezipped (Fig. 3e).

The results in Fig. 3 provide substantial evidence for RNA-DNA hybrid formation in front of a backtracked Pol II at a DNA fork in a head-on configuration. The formation of such a hybrid further anchors Pol II to the DNA substrate. It is important to note that dsDNA and RNA-DNA hybrid assume helical structures, although our cartoons did not show such helicity to ease strand tracking. Due to this helicity, unwinding the DNA leads to a concurrent rotation of the parental DNA and, consequently, the bound Pol II. This rotation can readily occur until RNA-DNA hybrid formation, which restricts Pol II rotation and DNA unwinding. Thus, RNA-DNA hybrid formation topologically locks Pol II on the DNA (Fig. 3d inset), making it difficult for the unzipping fork to remove the bound Pol II. This could explain why Pol II more strongly resists removal only in the presence of a long nascent RNA that enables the RNA-DNA hybrid formation in front of Pol II in the head-on configuration in Fig. 1e.

*In vivo*, this topological lock could be released if the 3’ RNA detaches from Pol II with the help of anti-backtracking factors, such as TFIIS, which facilitates Pol II cleavage of the 3’ RNA. The cleaved 3’ RNA segment can exit Pol II’s secondary channel, severing the connection between the Pol II and the RNA-DNA hybrid. To investigate this potential role of TFIIS, we performed similar experiments in the presence of TFIIS (without RNase T1) but with an additional step of attempting to unzip the DNA at the end to detect the final location of Pol II. For these experiments, we focused on traces that backtracked significantly in the force clamp step and could not be rezipped initially, consistent with RNA-DNA hybrid formation in front of Pol II.

We found that in the absence of TFIIS, the DNA remained unable to be rezipped, and Pol II had backtracked further when examined during the final step to attempt unzipping (Fig. 4b; Supplementary Fig. 5). This supports the possibility that the force on the DNA during the last two steps, even though small, could still promote Pol II backtracking. We observed this behavior even in the presence of TFIIS. However, with TFIIS, we also detected traces that showed a new behavior (Fig. 4c; Supplementary Fig. 5): even though a trace could not be rezipped initially, it subsequently became rezipped. In these traces, we found that Pol II regained its ability to translocate forward when we checked Pol II position during the last step. This behavior was never observed without TFIIS. The emergence of this new behavior is consistent with the interpretation that 3’ RNA cleavage of the RNA-DNA hybrid made it easier for the hybrid removal by either the DNA rezipping or by the forward translocation of Pol II.

**Figure 4.**
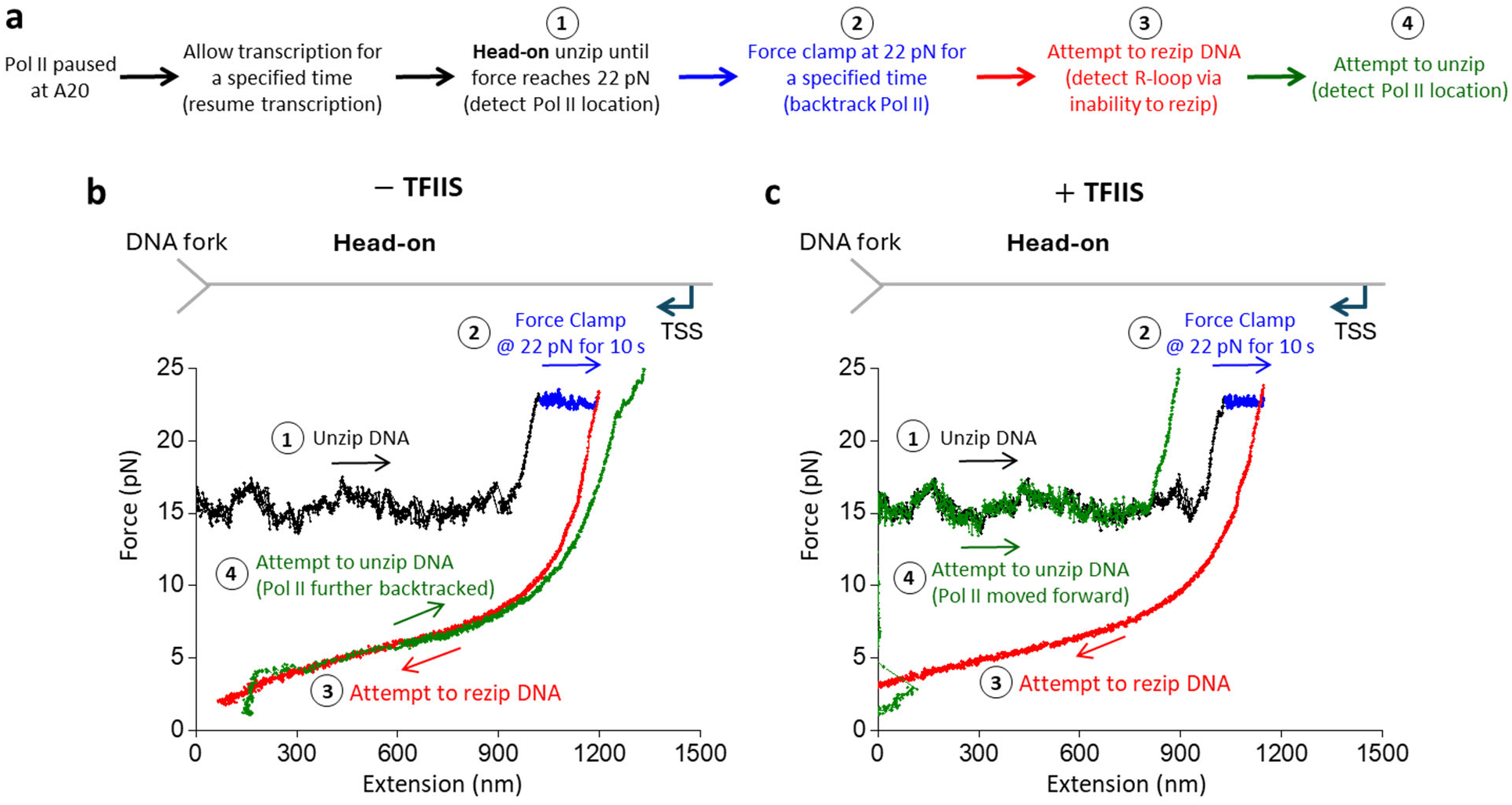
TFIIS facilitates RNA-DNA hybrid removal from backtracked Pol II. **a.** Outline of the experimental steps to backtrack Pol II, form RNA-DNA hybrid, and detect Pol II location after RNA-DNA hybridization. **b.** Representative force-extension trace showing the interaction of the DNA fork with an elongating Pol II in the head-on configuration without TFIIS. After Pol II was backtracked in step 2, DNA could not be rezipped in step 3. Pol II was further backtracked when checked during step 4. **c.** Representative force-extension trace showing the interaction of the DNA fork with an elongating Pol II in the head-on configuration with TFIIS present. After Pol II was backtracked in step 2, DNA could not be rezipped in step 3. However, DNA became rezipped and Pol II forward translocated when checked during step 4.

Collectively, our data may have significant *in vivo* implications for a head-on Pol II collision with a replisome. Replisome progression may backtrack Pol II, leading to RNA-DNA hybrid formation on the lagging strand in front of Pol II. This hybrid could topologically lock Pol II on DNA, exacerbating the Pol II roadblock to the replisome. TFIIS could facilitate Pol II cleavage of the 3’ RNA, which detaches the RNA-DNA hybrid from the bound Pol II and facilitates hybrid removal to alleviate the Pol II roadblock.

## RNA-DNA hybrid enables lagging-strand replication

*In vivo*, if an RNA-DNA hybrid forms on the lagging strand during a head-on collision of a replisome with Pol II, the RNA in this hybrid may serve as a primer for lagging-strand replication. This may alleviate some replication stress by efficiently leveraging the natural product in such a conflict, contributing to a mechanism for maintaining an active fork.

To explore this possibility, we extended our experimental approach used in Fig. 3 to enable lagging-strand replication (Fig. 5a). We first backtracked Pol II before attempting to rezip the DNA (Fig 5b). If the force continued to drop during the rezipping attempt, the inability to rezip indicated the formation of the RNA-DNA hybrid. We then allowed the force to decrease to around 1 pN before transitioning to a buffer containing T7 DNA polymerase and dNTPs that allow lagging-strand replication (Methods) (Fig. 5c,d). Under this force, one base pair of dsDNA has a longer extension than one nucleotide of ssDNA. This differential extension can then be used to indicate lagging-strand replication. Fig. 5c shows an example trace where the extension increased steadily with time, indicating steady lagging strand replication. The extension increase stopped at a position close to what would be expected, assuming all ssDNA on the lagging strand was converted to dsDNA (Fig. 5e). As a control, DNA extension changed minimally when the experiment was conducted without DNA polymerase (Fig. 5e).

**Figure 5.**
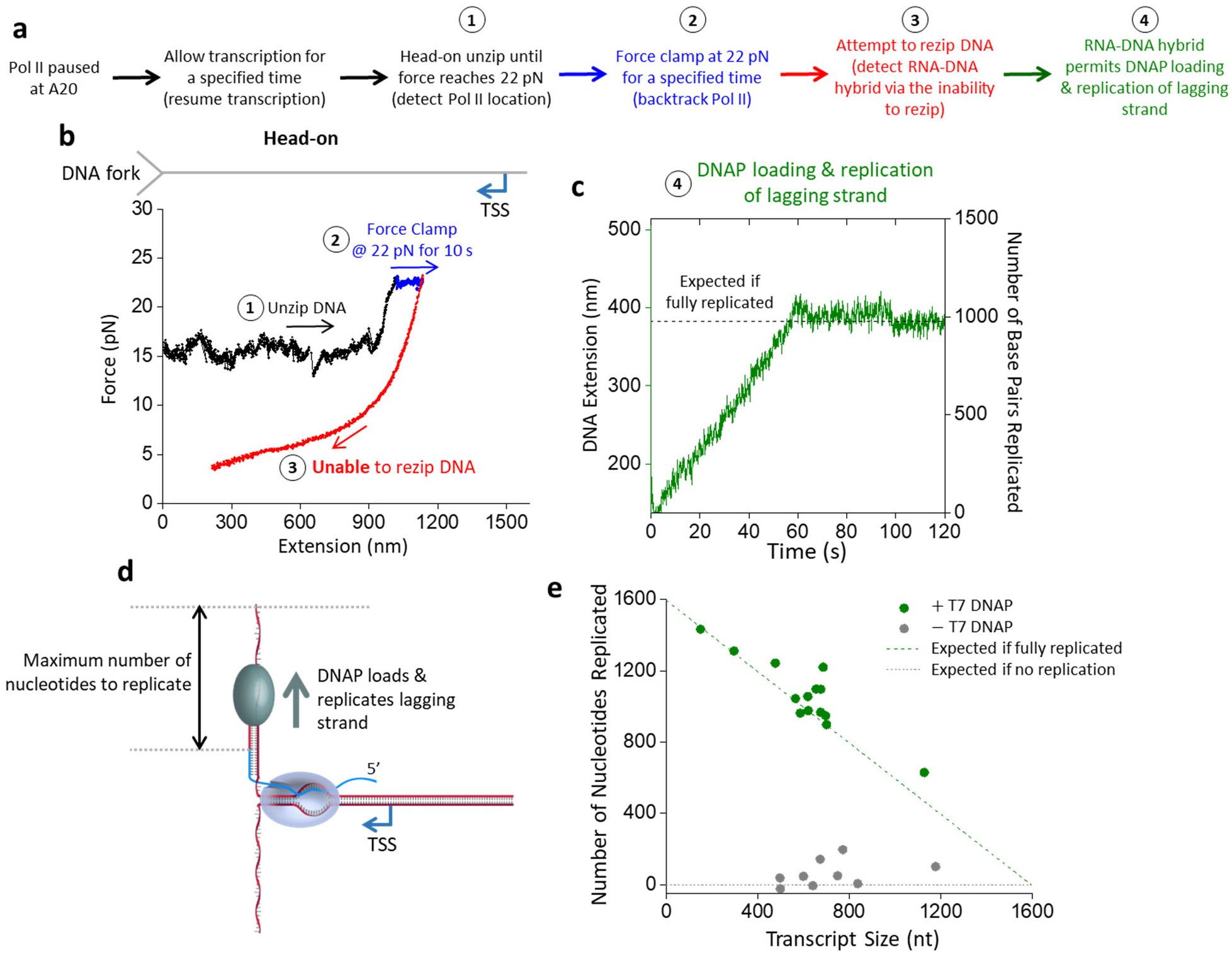
RNA-DNA hybrid formation enables lagging-strand replication. **a.** Experimental outline for steps to locate and backtrack Pol II, detect RNA-DNA hybrid formation, and observe T7 DNAP replication on the lagging strand. **b.** Representative force-extension trace of the DNA fork interaction with an elongating Pol II in the head-on configuration, with extensive backtracking. **c.** Monitoring the real-time DNA replication during step 4 of the trace shown in b. During this step, the force clamp was held at around 1 pN. Under this low force, DNA extension increased as T7 DNAP converted the ssDNA of the lagging strand into dsDNA. The right vertical axis shows the conversion from extension into the number of base pairs replicated. The black dashed line represents the position if T7 DNAP fully replicates the available lagging strand. **d.** Cartoon of T7 DNAP replication of the lagging strand using the RNA-DNA hybrid as a primer. **e.** Replication distance versus transcript size. The green dashed line indicates the expected number of nucleotides replicated if the lagging strand is fully replicated. The grey dashed line shows the expected number of nucleotides replicated without replication. (+) T7 DNAP, *N* = 14; (-) T7 DNAP, *N* = 9.

Therefore, we demonstrate that an RNA-DNA hybrid formed on the lagging strand during a head-on collision between a DNA fork and Pol II can enable lagging-strand replication. In a eukaryotic replisome, the lagging-strand replication is carried out by Pol δ, while we used the T7 DNA polymerase here for simplicity as a proof of principle. We hypothesize that Pol δ could perform a similar function during a replisome-transcription collision. Our proposed model does not require Pol II removal. Instead, it only requires Pol II to backtrack, which would be expected when Pol II collides with a replisome head-on. A backtracked Pol II then allows RNA-DNA hybrid formation in front of Pol II, and such a hybrid can then initiate lagging-strand replication even if Pol II remains bound.

## Discussion

In this work, we mimicked the replisome progression using mechanical unzipping of DNA. Using this approach, we investigated the consequences of an advancing DNA fork colliding with a transcribing Pol II either co-directionally or head-on. Our work provides a physical explanation for the polarity of the Pol II roadblock, demonstrates that an RNA-DNA hybrid can form not only behind Pol II but also in front of Pol II, and raises the possibility that the RNA-DNA hybrid formed in front of Pol II has the potential for initiation of lagging strand replication.

We show that Pol II roadblock polarity to the DNA fork is inherent to the Pol II elongation complex (Fig. 1): Pol II binds more stably to DNA and resists removal via sliding along the DNA in the head-on configuration, even when the transcript size is minimal. This intrinsic polarity could explain why transcription-replication conflicts are more likely to induce replisome stalling during a head-on transcription-replication conflict in the cell. Our unzipping mapper detected that Pol II minimally interacts with the DNA behind the transcription bubble but firmly clamps down on the DNA in front of the transcription bubble. Interestingly, we previously detected a similar interaction pattern for other complexes that also contain a DNA bubble with an RNA-DNA hybrid, such as the *E. coli* RNAP elongation complex^29,31,34,37^, a bound Cas9^31^, and a bound Cas12a^31^. The striking similarities among these complexes suggest a general design strategy among complexes containing an internal R-loop.

Although the Pol II roadblock has an inherent polarity, we found that the presence of a long RNA transcript strengthens Pol II’s interactions with DNA in both orientations while retaining the Pol II roadblock polarity (Fig. 1). We showed that an elongating Pol II with a long RNA transcript is also a more potent and persistent roadblock in the head-on configuration than in the co-directional configuration. We observed that a long RNA transcript allows RNA-DNA hybrid formation if there is available ssDNA near the Pol II (Fig. 2). Significantly, we made a surprising discovery that during a Pol II head-on collision with the DNA fork, an RNA-DNA hybrid can form on the lagging strand in front of Pol II after Pol II backtracks (Fig. 3). The enhanced Pol II interaction with DNA is likely a result of an RNA-DNA hybrid anchoring Pol II to the DNA via a topological lock (Fig. 3d). Consistent with this interpretation, the presence of TFIIS, which facilities Pol II cleavage of the 3’ RNA, allows RNA-DNA hybrid removal, suggesting that the connection between the bound Pol II and the RNA-DNA hybrid is severed. Interestingly, while the prevalent view places an RNA-DNA hybrid behind Pol II during a transcription-replication conflict^27^, recent studies showed that the presence of TFIIS is critical to maintaining genome stability and alluded to the possibility of the formation of an RNA-DNA hybrid in front of Pol II^52,53^. Our data now provide additional evidence of these emerging views.

Since the Pol II roadblock during a head-on collision threatens genome stability, it would be advantageous for the replisome to take advantage of this collision by using the RNA in the RNA-DNA hybrid in front of Pol II for lagging strand replication. We demonstrated the possibility of lagging strand replication using T7 DNAP. RNA-DNA hybrids have indeed been found on the lagging strand behind an eukaryotic replication fork in head-on collisions^25,54,55^. Whether these hybrids can serve as primers for the lagging strand replication *in vivo* remains to be seen.

Ultimately, the stalled Pol II must be moved off the DNA to allow replication progression^16^. It is possible that the replisome, especially with the help of other helicases, such as Pif1 and Rrm3^56–58^, is sufficiently powerful to evict the stalled Pol II. Factors directly interacting with Pol II could also assist Pol II eviction, but they are yet to be identified. Although Mfd in prokaryotes can evict an RNAP stalled at a DNA fork^16,29,59–61^, its eukaryotic counterparts, CSB (Human) or Rad26 (S. cerevisiae), do not exhibit this capability ^62,63^.

We hypothesize that an RNA-DNA hybrid can form in front of Pol II only after Pol II physically encounters the replication fork during a head-on transcription-replication conflict. However, when Pol II approaches a replisome head-on, (+) torsion accumulates between Pol II and the replisome well before Pol II physically encounters the replisome, because torsion in the DNA can act over a distance^23^ .This (+) torsion accumulation could induce replication fork stalling and disassembly as well as Poll II backtracking and stalling^18,23^. Topoisomerases can resolve torsional stress during these conflicts, but their impact on the RNA-DNA hybrid formation during these conflicts is a double-edged sword. Before Pol II encounters a replisome, supercoiling relaxation allows Pol II progression and limits its backtracking, potentially reducing the RNA-DNA hybrid formation at the fork once the collision occurs. However, supercoiling relaxation increases the opportunity for collision by allowing the replisome and Pol II to come into direct contact, which then permits RNA-DNA hybrid formation at the fork. Previous studies of the head-on transcription-replication conflict found that supercoiling resolution by gyrase in prokaryotes drives RNA-DNA hybrid formation^18^, whereas supercoiling resolution by topoisomerase I in eukaryotes prevents RNA-DNA hybrid formation^4,44,64,65^.

Our work uses a DNA fork to mimic the replisome and may not capture the full complexity of what might occur during transcription-replication conflicts *in vivo*. However, the simplicity of our model system permits mechanistic studies and reveals important physical parameters for the formation of the RNA-DNA hybrid. We now have a significantly clearer understanding of the nature of the RNA-DNA hybrid. We anticipate what we have learned from this work may facilitate data interpretation of more complex *in vivo* systems.

## ACKNOWLEDGEMENTS

We thank members of the Michelle Wang Laboratory for helpful discussion and comments. We also thank Juntaek Oh and Peini Hou for purifying the 10-subunit Pol II used for initial troubleshooting of the EC assembly. This work is supported by the National Institutes of Health grants R01GM136894 (to M.D.W.), T32GM008267 (to M.D.W.), and GM102362 (to D.W.). M.D.W. is a Howard Hughes Medical Institute investigator.

## AUTHOR CONTRIBUTIONS

T.M.K., J.T.I, T.T.L, and M.D.W. designed single-molecule assays. T.M.K., J.Q., and P.M.H designed DNA substrates. T.M.K. prepared DNA substrates. L.L. and M.K. purified and characterized the 12-subunit Pol II and TFIIS. T.M.K. performed single-molecule experiments. J.Q. and P.M.H provided technical advice. T.M.K. and J.T.I. analyzed data. T.T.L. performed some preliminary experiments using E. coli RNA polymerase. D.W. provided the 10-subunit Pol II used only for initial troubleshooting of the EC assembly. M.D.W. wrote the initial draft. All authors contributed to the manuscript revision. M.D.W. supervised this project.

## DATA AND MATERIAL AVAILABILITY

All data are available in the main text or the supplementary materials.

## References

1 Deshpande, A. M. & Newlon, C. S. DNA Replication Fork Pause Sites Dependent on Transcription. Science 272, 1030–1033 (1996).

2 Takeuchi, Y., Horiuchi, T. & Kobayashi, T. Transcription-dependent recombination and the role of fork collision in yeast rDNA. Genes Dev 17, 1497–1506 (2003).

3 Azvolinsky, A., Giresi, P. G., Lieb, J. D. & Zakian, V. A. Highly transcribed RNA polymerase II genes are impediments to replication fork progression in Saccharomyces cerevisiae. Mol Cell 34, 722–734 (2009).

4 Tuduri, S. et al. Topoisomerase I suppresses genomic instability by preventing interference between replication and transcription. Nat Cell Biol 11, 1315–1324 (2009).

5 Helmrich, A., Ballarino, M. & Tora, L. Collisions between Replication and Transcription Complexes Cause Common Fragile Site Instability at the Longest Human Genes. Molecular Cell 44, 966–977 (2011).

6 Martin, M. M. et al. Genome-wide depletion of replication initiation events in highly transcribed regions. Genome Res 21, 1822–1832 (2011).

7 Wang, J. et al. Persistence of RNA transcription during DNA replication delays duplication of transcription start sites until G2/M. Cell Reports 34, 108759 (2021).

8 St Germain, C. P., et al. Genomic patterns of transcription-replication interactions in mouse primary B cells. Nucleic Acids Res 50, 2051–2073 (2022).

9 Browning, K. R. & Merrikh, H. Replication-Transcription Conflicts: A Perpetual War on the Chromosome. Annu Rev Biochem 93, 21–46 (2024).

10 French, S. Consequences of replication fork movement through transcription units in vivo. Science 258, 1362–1365 (1992).

11 Prado, F. & Aguilera, A. Impairment of replication fork progression mediates RNA polII transcription-associated recombination. EMBO J 24, 1267–1276 (2005).

12 Mirkin, E. V. & Mirkin, S. M. Mechanisms of transcription-replication collisions in bacteria. Mol Cell Biol 25, 888–895 (2005).

13 Liu, B. & Alberts, B. M. Head-on collision between a DNA replication apparatus and RNA polymerase transcription complex. Science 267, 1131–1137 (1995).

14 Kim, N., Abdulovic, A. L., Gealy, R., Lippert, M. J. & Jinks-Robertson, S. Transcription-associated mutagenesis in yeast is directly proportional to the level of gene expression and influenced by the direction of DNA replication. DNA Repair 6, 1285–1296 (2007).

15 Pomerantz, R. T. & O’Donnell, M. The replisome uses mRNA as a primer after colliding with RNA polymerase. Nature 456, 762–766 (2008).

16 Pomerantz, R. T. & O’Donnell, M. Direct restart of a replication fork stalled by a head-on RNA polymerase. Science 327, 590–592 (2010).

17 Merrikh, C. N., Brewer, B. J. & Merrikh, H. The B. subtilis Accessory Helicase PcrA Facilitates DNA Replication through Transcription Units. PLoS Genet 11 (2015).

18 Lang, K. S. & Merrikh, H. Topological stress is responsible for the detrimental outcomes of head-on replication-transcription conflicts. Cell Rep 34, 108797 (2021).

19 Paul, S., Million-Weaver, S., Chattopadhyay, S., Sokurenko, E. & Merrikh, H. Accelerated gene evolution through replication–transcription conflicts. Nature 495, 512–515 (2013).

20 Gan, W. et al. R-loop-mediated genomic instability is caused by impairment of replication fork progression. Genes Dev 25, 2041–2056 (2011).

21 Santos-Pereira, J. M. & Aguilera, A. R loops: new modulators of genome dynamics and function. Nature Reviews Genetics 16, 583–597 (2015).

22 Merrikh, H., Machón, C., Grainger, W. H., Grossman, A. D. & Soultanas, P. Co-directional replication-transcription conflicts lead to replication restart. Nature 470, 554–557 (2011).

23 Lang, K. S. et al. Replication-Transcription Conflicts Generate R-Loops that Orchestrate Bacterial Stress Survival and Pathogenesis. Cell 170, 787–799 (2017).

24 Hamperl, S., Bocek, M. J., Saldivar, J. C., Swigut, T. & Cimprich, K. A. Transcription-Replication Conflict Orientation Modulates R-Loop Levels and Activates Distinct DNA Damage Responses. Cell 170, 774–786 e719 (2017).

25 Kumar, C. & Remus, D. Looping out of control: R-loops in transcription-replication conflict. Chromosoma 133, 37–56 (2024).

26 Roy, D., Yu, K. & Lieber, M. R. Mechanism of R-Loop Formation at Immunoglobulin Class Switch Sequences. Molecular and Cellular Biology 28, 50–60 (2008).

27 Hamperl, S. & Cimprich, K. A. The contribution of co-transcriptional RNA:DNA hybrid structures to DNA damage and genome instability. DNA Repair (Amst) 19, 84–94 (2014).

28 Le, T. T. et al. Etoposide promotes DNA loop trapping and barrier formation by topoisomerase II. Nat Chem Biol 19, 641–650 (2023).

29 Le, T. T. et al. Mfd Dynamically Regulates Transcription via a Release and Catch-Up Mechanism. Cell 172, 344–357.e315 (2018).

30 Ye, F., Inman, J. T., Hong, Y., Hall, P. M. & Wang, M. D. Resonator nanophotonic standing-wave array trap for single-molecule manipulation and measurement. Nat Commun 13, 77 (2022).

31 Hall, P. M. et al. Polarity of the CRISPR roadblock to transcription. Nat Struct Mol Biol 29, 1217–1227 (2022).

32 Sun, B. et al. Helicase promotes replication re-initiation from an RNA transcript. Nat Commun 9, 2306 (2018).

33 Walter, W., Kireeva, M. L., Studitsky, V. M. & Kashlev, M. Bacterial Polymerase and Yeast Polymerase II Use Similar Mechanisms for Transcription through Nucleosomes*. Journal of Biological Chemistry 278, 36148–36156 (2003).

34 Inman, J. T. et al. DNA y structure: a versatile, multidimensional single molecule assay. Nano Lett 14, 6475–6480 (2014).

35 Kireeva, M. L., Komissarova, N., Waugh, D. S. & Kashlev, M. The 8-Nucleotide-long RNA:DNA Hybrid Is a Primary Stability Determinant of the RNA Polymerase II Elongation Complex*. Journal of Biological Chemistry 275, 6530–6536 (2000).

36 Komissarova, N. & Kashlev, M. Functional topography of nascent RNA in elongation intermediates of RNA polymerase. Proc Natl Acad Sci U S A 95, 14699–14704 (1998).

37 Jin, J. et al. Synergistic action of RNA polymerases in overcoming the nucleosomal barrier. Nat Struct Mol Biol 17, 745–752 (2010).

38 Cramer, P. et al. Architecture of RNA Polymerase II and Implications for the Transcription Mechanism. Science 288, 640–649 (2000).

39 Armache, K.-J., Kettenberger, H. & Cramer, P. Architecture of initiation-competent 12-subunit RNA polymerase II. Proceedings of the National Academy of Sciences 100, 6964–6968 (2003).

40 Bushnell, D. A. & Kornberg, R. D. Complete, 12-subunit RNA polymerase II at 4.1-Å resolution: Implications for the initiation of transcription. Proceedings of the National Academy of Sciences 100, 6969–6973 (2003).

41 Drolet, M., Bi, X. & Liu, L. F. Hypernegative supercoiling of the DNA template during transcription elongation in vitro. J Biol Chem 269, 2068–2074 (1994).

42 Drolet, M. et al. Overexpression of RNase H partially complements the growth defect of an Escherichia coli delta topA mutant: R-loop formation is a major problem in the absence of DNA topoisomerase I. Proc Natl Acad Sci U S A 92, 3526–3530 (1995).

43 Drolet, M. et al. The problem of hypernegative supercoiling and R-loop formation in transcription. Front Biosci 1, 970 (2003).

44 El Hage, A., French, S. L., Beyer, A. L. & Tollervey, D. Loss of Topoisomerase I leads to R-loop-mediated transcriptional blocks during ribosomal RNA synthesis. Genes Dev 24, 1546–1558 (2010).

45 Komissarova, N. & Kashlev, M. Transcriptional arrest: *Escherichia coli* RNA polymerase translocates backward, leaving the 3&#x2032; end of the RNA intact and&#x2009;extruded. Proceedings of the National Academy of Sciences 94, 1755-1760 (1997).

46 Wang, D. et al. Structural basis of transcription: backtracked RNA polymerase II at 3.4 angstrom resolution. Science 324, 1203–1206 (2009).

47 Cheung, A. C. & Cramer, P. Structural basis of RNA polymerase II backtracking, arrest and reactivation. Nature 471, 249–253 (2011).

48 Nudler, E. RNA polymerase backtracking in gene regulation and genome instability. Cell 149, 1438–1445 (2012).

49 Burgers, P. M. J. & Kunkel, T. A. Eukaryotic DNA Replication Fork. Annu Rev Biochem 86, 417–438 (2017).

50 Pike, A. M., Friend, C. M. & Bell, S. P. Distinct RPA functions promote eukaryotic DNA replication initiation and elongation. Nucleic Acids Research 51, 10506–10518 (2023).

51 Jones, M. L., Aria, V., Baris, Y. & Yeeles, J. T. P. How Pol α-primase is targeted to replisomes to prime eukaryotic DNA replication. Mol Cell 83, 2911–2924 (2023).

52 Zatreanu, D. et al. Elongation Factor TFIIS Prevents Transcription Stress and R-Loop Accumulation to Maintain Genome Stability. Mol Cell 76, 57–69 e59 (2019).

53 Duardo, R. C. et al. Human DNA topoisomerase I poisoning causes R loop-mediated genome instability attenuated by transcription factor IIS. Sci Adv 10, eadm8196 (2024).

54 Stoy, H. et al. Direct visualization of transcription-replication conflicts reveals post-replicative DNA:RNA hybrids. Nat Struct Mol Biol 30, 348–359 (2023).

55 Kumar, C., Batra, S., Griffith, J. D. & Remus, D. The interplay of RNA:DNA hybrid structure and G-quadruplexes determines the outcome of R-loop-replisome collisions. Elife 10, e72286 (2021).

56 Ivessa, A. S. et al. The Saccharomyces cerevisiae Helicase Rrm3p Facilitates Replication Past Nonhistone Protein-DNA Complexes. Molecular Cell 12, 1525–1536 (2003).

57. Tran, P. L. T., et al. PIF1 family DNA helicases suppress R-loop mediated genome instability at tRNA genes. Nat Commun 8 (2017).

58 Torres, J. Z., Bessler, J. B. & Zakian, V. A. Local chromatin structure at the ribosomal DNA causes replication fork pausing and genome instability in the absence of the S. cerevisiae DNA helicase Rrm3p. Genes Dev 18, 498–503 (2004).

59 Selby, C. P. & Sancar, A. Molecular Mechanism of Transcription-Repair Coupling. Science 260, 53–58 (1993).

60 Park, J.-S., Marr, M. T. & Roberts, J. W. E. coli Transcription Repair Coupling Factor (Mfd Protein) Rescues Arrested Complexes by Promoting Forward Translocation. Cell 109, 757–767 (2002).

61 Howan, K. et al. Initiation of transcription-coupled repair characterized at single-molecule resolution. Nature 490, 431–434 (2012).

62 Selby, C. P. & Sancar, A. Human transcription-repair coupling factor CSB/ERCC6 is a DNA-stimulated ATPase but is not a helicase and does not disrupt the ternary transcription complex of stalled RNA polymerase II. Journal of Biological Chemistry 272, 1885–1890 (1997).

63 Xu, J. et al. Structural basis for the initiation of eukaryotic transcription-coupled DNA repair. Nature 551, 653–657 (2017).

64 Yang, Y. et al. Arginine Methylation Facilitates the Recruitment of TOP3B to Chromatin to Prevent R Loop Accumulation. Molecular Cell 53, 484–497 (2014).

65 Pommier, Y., Sun, Y., Huang, S.-y. N. & Nitiss, J. L. Roles of eukaryotic topoisomerases in transcription, replication and genomic stability. Nature Reviews Molecular Cell Biology 17, 703–721 (2016).

