## Supplementary Materials for "RNA Polymerase II is a Polar Roadblock to a Progressing DNA Fork"

### Materials and Methods

#### Protein expression and purification

Yeast RNA Polymerase II (Pol II) containing a 6xHis-tag on both Rpb1 and Rpb3 subunits was expressed and purified from *S. cerevisiae*[1]. In brief, cell pellets were lysed and applied to a HisTrap HP column (Cytiva 17524801), a heparin column (Cytiva 17040703) and a Mono Q column 10/100 (Cytiva 17516701). The eluted Pol II was assessed via SDS-PAGE 4-12% Bis-Tris SDS gels in 1X MES SDS buffer, then concentrated and flash-frozen for storage.

*S. cerevisiae* recombinant TFIIS was expressed in *BL21 (DE3) pET15b PPR1* (69450-M; Novagen-sigma)[2,3]. Cell pellets were harvested by centrifugation and then sonicated using a microprobe tip and Sonicator-Ultrasonic Processor VHX 750 watt (model GEX 750; PG Scientific). Each chromatography step was assessed via SDS-PAGE with 12% Bis-Tris SDS gels in 1X MES SDS buffer. The centrifugated lysate was applied to a HisTrap FF Crude column (Cytiva 17528601) and then applied to a Mono S column 10/100 GL (Cytiva 17516901). The eluted protein eluted was diluted and applied to a HisTrap HP (Cytiva 29051021) column and a Mono S column 5/50 GL (Cytiva 17516801) to produce 6xHis-tagged protein. The protein was assessed via SDS-PAGE, concentrated, and flash-frozen for storage.

Wild-type T7 DNA polymerase was purchased from New England Biolabs (NEB, M0274S). *E. coli* thioredoxin (trx) was purchased from Sigma-Aldrich (Sigma, T0910). RNase T1 was purchased from Thermo Fisher Scientific (Thermo Fisher, EN0542).

#### DNA Substrates

We utilized a Y-shaped DNA substrate mimicking a DNA replication fork[4-10] (Supplementary Fig. 2) that enabled DNA unzipping of a parental strand with Pol II assembled in the elongation complex. This Y-shaped DNA structure consists of Y-arms (fork-like structure with two daughter strands) and a parental DNA strand with assembled Pol II EC.

The Y-arms consist of both leading and lagging daughter strands. Two 4.15 kb daughter strands of identical sequence were amplified via PCR from pBR322 (NEB N3033S). The leading strand was amplified with a forward primer containing a 5’ digoxigenin (dig) label, while the lagging strand was amplified with a forward primer containing a 5’ biotin (bio) label (Supplementary Table 1). Each resulting arm was digested with BsmBi-V2 (NEB R0739S) and ligated with T4 DNA Ligase (NEB M0202) to the respective leading (Upper Leading Strand and Lower Leading Strand) or lagging (Upper Lagging Strand and Lower Lagging Strand) strand annealed adapter oligo (Supplementary Table 1). The “lower” leading and lagging adapter oligos contain a 30 nt sequence of complimentary ssDNA allowing the two pieces to be annealed to one another and create the Y-arms.

Figures 1-4 used a Y-arm structure with no additional modifications (Upper Leading Strand), while Figure 5 had a modified leading strand adapter oligo (Upper Leading Strand_invdT) with an inverted dT on the 3' end to prevent T7 DNAP from loading and replicating the leading strand.

The complete Y-shaped DNA substrate was created by ligation of the Y-arms to a parental DNA strand via a unique 3' overhang. Two parental strands were generated for this study: co-directional (CD) or head-on (HO). Both strands consisted of three key pieces: the transcription elongation complex (TEC), a downstream segment, and an upstream segment.

Both templates used an identical TEC DNA segment, consisting of four annealed complimentary ssDNA oligos assembled with RNA Polymerase II[11-13]. Annealing resulted in a short 130 bp DNA construct with two unique overhangs to allow for ligation to the downstream and upstream templates. A downstream non-template strand (DS NTS), an upstream NTS (US NTS), and a template strand (TS) (Supplementary Table 1) were first annealed to create a 'gapped' DNA construct, with a 76 nt gap exposing the TS as ssDNA between the two NTS pieces. A 14 nt RNA (RNA 14), with 9 nt complimentary to the TS, was annealed to the gapped region, forming an RNA-DNA hybrid scaffold. Pol II was added to the hybrid and incubated at room temperature before the addition of a complementary 76 nt NTS ssDNA oligo (ARP77). The nicks at the two ends of the gap were ligated by T4 DNA Ligase (M0202) in the same reaction listed below that ligates the downstream, upstream, and Y-arm structure together.

For the CD parental strand, the upstream segment was amplified from pBR322 (NEB N3033S) and enzymatically digested with both PpuMI (NEB R0506) and DraIII-HF (NEB R3510S) to generate a 0.65 kb strand. The downstream segment was amplified from pRL574 and digested with BstXI (NEB R0113), generating a 1.5 kb strand. The pRL574 plasmid, containing the partial beta subunit gene, was used to allow for a readable transcription sequence, but the T7 A1 promoter and the T7 TSS were not in the amplified region to avoid sequence overlap with the Pol II TEC sequence. The upstream segment was ligated to the Y-arms and the upstream end of the TEC, and the downstream segment was ligated to the downstream end of the TEC.

For the HO parental strand, which was assembled in the opposite direction, the upstream segment was amplified from pBR322 (NEB) but only digested with PpuMI (NEB R0506), generating a 1.4 kb strand. The downstream template was amplified from the same region of pRL574 as the CD template, but digested with both BstXI (NEB R0113) and AlwNI (NEB R0514S), generating a 1.5 kb strand. The upstream segment was ligated to the upstream end of the TEC and the downstream segment was ligated to the Y-arms and the downstream side of the TEC.

For each CD or HO template, the Y-arm structure, appropriate corresponding downstream and upstream segments, and the TEC were ligated together using T4 DNA Ligase (NEB M0202) in the same reaction.

**Experimental Conditions**

All experiments were conducted in a multi-channel laminar flow cell at room temperature (23 C). Figures 1-4 used a four-channel flow cell, while Figure 5 used a five-channel flow cell.

In the four-channel flow cell, Channel 1 contained anti-digoxigenin coated polystyrene beads (1 um, Polysciences, 08226-15) pre-incubated with a DNA Y-structure template. Beads were suspended in a 1x Transcription Buffer (TB150) (25 mM Tris-HCl pH 8.0, 150 mM KCl, 10 µM ZnSO4, 1 mM DTT, 2 mM TCEP, 3% Glycerol) with 1 mM $MgCl_2$. Channel 2 contained A20 Buffer (1x TB150 with 4 mM $MgCl_2$, 1 mM ATP, 1 mM GTP, and 1 mM CTP) for Pol II elongation to an Adenine (A) 20 nt away from the TSS. Channel 3 contained NTP Buffer (1X TB150 with 5 mM MgCl2, and 1 mM of each NTP) for full transcription. Both channel 2 and 3 contained an oxygen scavenger system (10 nM protocatechuate-3,4-dioxygenase and 2.5 mM protocatechuic acid) to minimize photodamage to the DNA and increase tether lifetime. Channel 4 contained streptavidin coated polystyrene beads (1 um, Polysciences, 08226-15) in 1x PBS (137 mM NaCl, 2.7 mM KCl, 8 mM Na2 HPO4, and 2 mM KH2PO4).

Variations in Channels 2 and 3 were present for different experimental conditions. In Figures 1-3, (-) RNase conditions had 0.25 u/µL of SUPERase·In RNase Inhibitor (Thermo Fisher AM2696). For data taken in a (+) RNase environment, channel 3 contained 10 u/µL of RNase T1 (Thermo Fisher, EN0541). In Figure 4, for data taken in the (+) TFIIS environment, Channel 3 was preincubated with 0.5 mg/mL acetylated BSA in 1x PBS, then exchanged to NTP buffer with 1 µM of wt TFIIS.

In the five-channel flow cell, Channels 1-3 were identical to those in the four-channel flow cell, with Channels 2 and 3 containing both the oxygen scavenger system and 0.25 u/µL of SUPERase·In RNase Inhibitor. Channel 4 contained 1x replication buffer (RB) (50 mM Tris-HCl (pH 7.5), 40 mM NaCl, 1.5 mM EDTA, 8 mM MgCl2, 1 mM DTT, 2 mM TCEP, and 3% glycerol) with 1 mM of each dNTP and 0.5 mg/mL β-casein, along with the oxygen scavenger system. In the presence of T7 DNAP, 20 nM wt T7 gp5 DNA polymerase and 150 nM thioredoxin were included. Channel 5 contained streptavidin coated polystyrene beads (1 µm) in 1x PBS.

**Single-molecule optical tweezers assay**

Data was collected using a home-built dual optical trap in combination with a microfluidic multi-channel laminar flow cell[4,14]. Each channel of the flow cell was fed with a 1 mL glass syringe (Hamilton, 81320), driven by a syringe pump with constant flow of 1.5 µL/min. The four-channel flow cell had a flow rate of 170 µm/s and the five-channel flow cell had a flow rate of 210 µm/s at the trapping position.

The DNA template was tethered between two optically trapped beads via its labeled daughter strands. A strep-coated polystyrene bead was trapped in a steered trap and tethered via the 5' biotin label on the lagging daughter strand, while an anti-dig-coated bead, pre-incubated with DNA, was tethered via the 5' dig label on the leading strand. Tether formation occurred in channel 2, allowing Pol II to escape from the TEC to an A20 pause site. The tether was then transferred to channel 3 to begin data collection.

**Single-molecule experimental procedures**

All experiments began by mechanically unzipping the parental DNA at 100 nm/s to mimic the replication fork. Figures 1 and 2 unzipped until the end of the template was reached, thereby unzipping through and probing the fork interaction with the Pol II. Figures 3-5 unzipped the duplex DNA until the Pol II molecule was encountered. This was determined by the force reaching 7 pN above the naked DNA baseline. A constant force (7 pN above the DNA baseline) was applied for 10 seconds to induce backtracking. The ability to and the extent of backtracking varied between molecules. Figures 3-5 tested for RNA-DNA hybrid formation by an inability to reanneal the duplex DNA in front of the Pol II after backtracking. This was tested using by mechanically rezipping the DNA at a velocity of -100 nm/s. In Figure 5, after an RNA-DNA hybrid was identified, tethers were moved into Channel 4 and held at a constant force of around 1 pN for 2 minutes to test for T7 DNAP replication of the lagging strand.

**Data acquisition and data conversion**

Data was acquired at 10 kHz and converted into force and extension as previously described[15]. All extension curves were modified to exclude the extension component contributed by the dsDNA arms using the worm-like chain theory. Elasticity parameters for dsDNA were obtained previously[16], and ssDNA and RNA-DNA hybrids were measured in lab using force-extension curves. Due to bead size variations, the trap stiffness of each bead pair was calibrated using the theoretically predicted force-extension curve and the measured unzipping force preceding the DNA fork's encounter with a bound Pol II.

**Unzipping Alignment**

To improve the precision and accuracy in the unzipping data, the extension of each bead pair was aligned to the theoretically predicted force-extension curve, using regions of the unzipping data that preceded the encounter of the DNA fork with Pol II.

**Single-molecule Data Analysis**

The transcript size of the nascent RNA was calculated using the position of the first encounter with Pol II by the DNA fork and the known TSS. Deviations in the force-extension curve from the theoretical naked DNA curve were used to indicate the Pol II location. These deviations could be seen as force dips, rises or extension shortening events, indicating a protein bound to the DNA substrate. In the case of a co-directional elongating Pol II in a (+) RNase condition, Pol II location was identified using a rezipping step to identify the location along the template at which the duplex could not reanneal. The extension at which Pol II was located was then converted into the number of base pairs unzipped, using the freely join chain model (FJC), allowing the transcript size to be identified.

The maximum displacement force was determined by measuring the maximum force reached while the fork interacted with Pol II, before complete disruption. The sliding distance was the total distance required for the fork to disrupt the Pol II-DNA interaction. Disruption of the Pol II-dsDNA interaction were determined by the return of the force-extension curve to the theoretical DNA force baseline.

RNA-DNA hybrid formation size was measured by analyzing the shift in the number of unzipped base pairs from the predicted force-unzipped base pairs curve for the given DNA sequence. After the first encounter with Pol II, a shift was applied to align the data with the DNA baseline. For co-directional traces this shift was aligned directly after Pol II disruption, while the head-on was shifted after the TSS site to ensure only the upstream DNA segment was being aligned. This shift was then used to calculate the RNA-DNA hybrid size using the FJC model for the ssDNA component and the WLC model for the RNA-DNA hybrid component.

The Pol II backtrack distance was calculated by subtracting the initial Pol II location from its new location after the 10 second ~22 pN constant force clamp step.

T7 DNAP replication was measured by the extension change that occurred during the ~1 pN constant force clamp step. The overall extension was adjusted by subtracting off the extension contributions from the leading strand, Pol II backtrack distance, and RNA-DNA hybrid components, leaving the extension of the lagging strand available to be replicated. Due to secondary structures in the ssDNA[15], the FJC model cannot be used for converting ssDNA into nucleotides at a low force. Instead, a conversion factor (28 nt/nm) was measured in a (-) T7 DNAP environment around a force of 1 pN. The lagging strand extension was then converted into the number of nucleotides replicated using the WLC for the dsDNA and the measured conversion factor for the ssDNA.

**Quantification and statistical analysis**

All data were obtained from at least eight independent replicates. Statistical details of individual experiments, including number of traces, mean, and SEM values can be found in the manuscript text, figures, and figure legends.

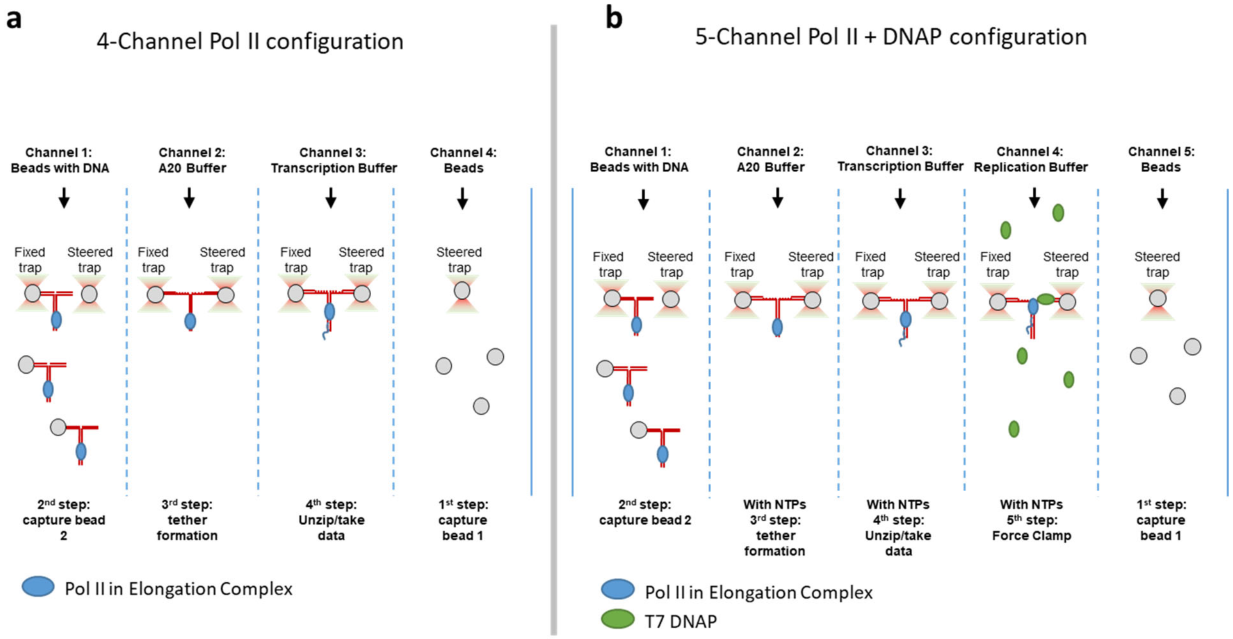


**Supplementary Figure 1.** The multi-channel microfluidic flow cells
**a.** The four-channel flow cell used for data collection for Figures 1, 2, 3, and 4. This flow cell, combined with a dual-trap optical tweezer, utilized laminar flow to partition different buffers, allowing for tether formation and movement into various experimental conditions. This allowed for transcription to only begin after a tether was formed and ready for data collection.
**b.** The five-channel flow cell used for data collection for Figure 5. This flow cell, combined with a dual-trap optical tweezer, utilized laminar flow to partition different buffers, allowing for tether formation and movement into various experimental conditions. This allowed for a single Pol II molecule to begin transcription and then be moved into a replication channel only after RNA-DNA hybrid formation.

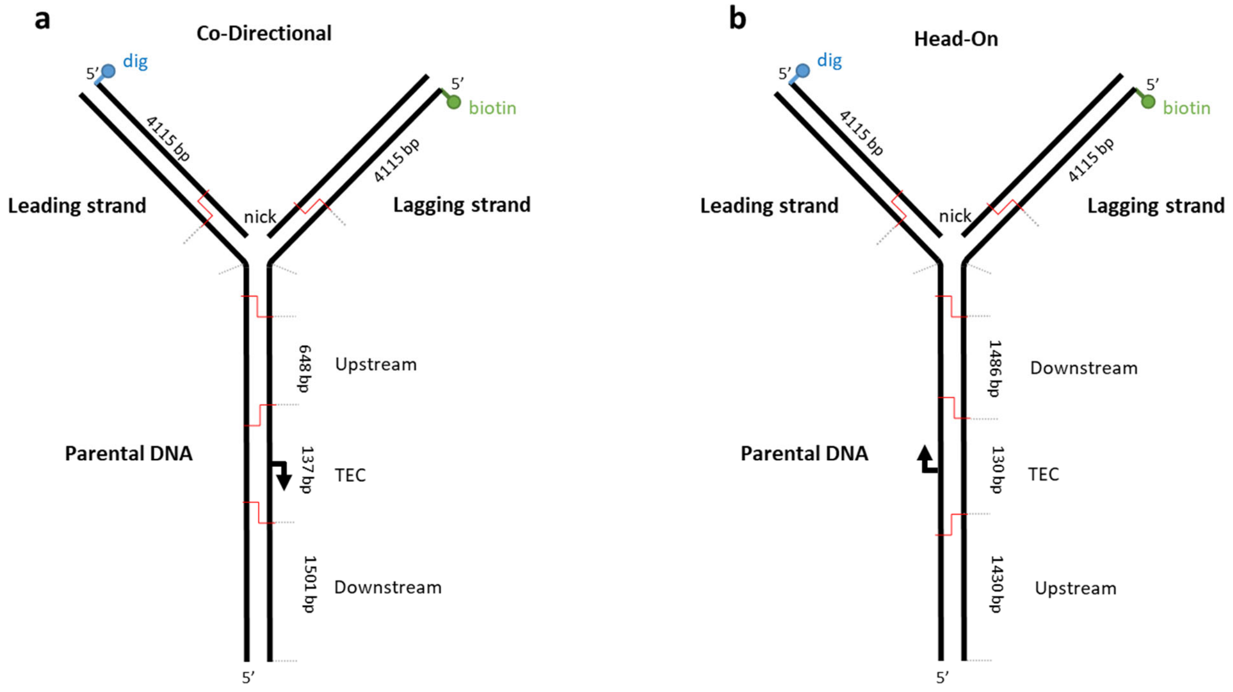


**Supplementary Figure 2.** The Y-shaped DNA substrates

**a.** The co-directional Y-shaped DNA substrate. The template consists of two labeled daughter strands to allow for tethering between the trapped beads. The daughter strands are connected to a three-way junction to resemble a replication fork. The parental strand is assembled from the fork as follows: upstream DNA, assembled Pol II TEC, and then downstream DNA, allowing for co-directional collision studies where Pol II moves away from the DNA fork.

**b.** The head-on Y-shaped DNA substrate. The template consists of two labeled daughter strands to allow for tethering between the trapped beads. The daughter strands are connected to a three-way junction to resemble a replication fork. The parental strand is assembled from the fork as follows: downstream DNA, assembled Pol II TEC, and then upstream DNA, allowing for head-on collision studies where Pol II moves towards the DNA fork.

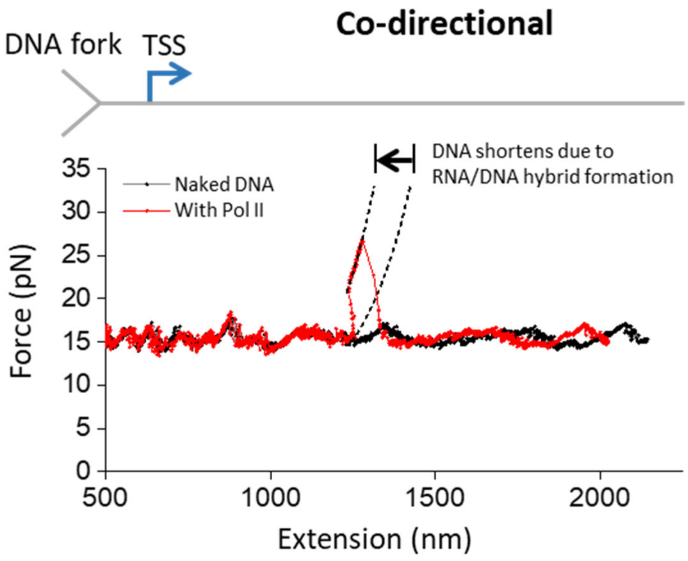


**Supplementary Figure 3.** Co-directional RNA-DNA hybrid extension shift in force peak Representative trace showing the interaction of the DNA fork with an elongating Pol II elongation complex (EC) in the co-directional (CD) orientation. The force-extension data of DNA with Pol II bound are shown in red while those of naked DNA of the same sequence is shown in black. As shown in Figure 2a, immediately upon the unzipping fork encountering the bound Pol II from behind, the force-extension curve snapped back and followed a curve at a shorter extension until the disruption of the bound Pol II. The Two dashed curves are used to show the initial Pol II position (right) and the Pol II position after RNA-DNA hybrid formation (left) before disruption of the protein. This snap-back behavior may be a result of RNA-DNA hybrid formation behind Pol II, with RNA-DNA hybrid formation on the leading strand shortening the overall DNA extension (Fig. 2b). The initial Pol II position is shown using a freely-jointed chain (FJC) model, with only ssDNA contributing; the shifted Pol II position is plotted using a combined FJC and worm-like chain (WLC) model due to having a hybrid component present. The black arrow indicates the amount of extension shortening. The TSS is illustrated in the cartoon above, with the blue arrow indicating the direction of movement of elongating Pol II.

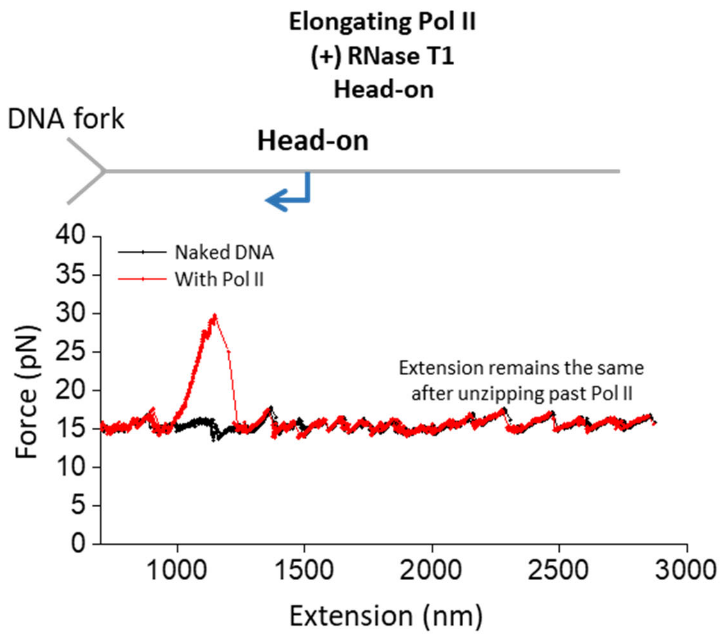


**Supplementary Figure 4.** Head-on RNase T1 example trace

Representative trace showing the interaction of the DNA fork with an elongating Pol II elongation complex (EC) in the head-on (HO) orientation in a (+) RNase T1 environment. The force-extension data of DNA with Pol II bound are shown in red while those of naked DNA of the same sequence are shown in black. No shift in extension is measured after disruption of Pol II or unzipping past the TSS, due to a lack of RNA-DNA hybrid formation as the majority of the RNA should have been digested by RNase T1. The TSS is illustrated in the cartoon above, with the blue arrow indicating the direction of elongating Pol II.

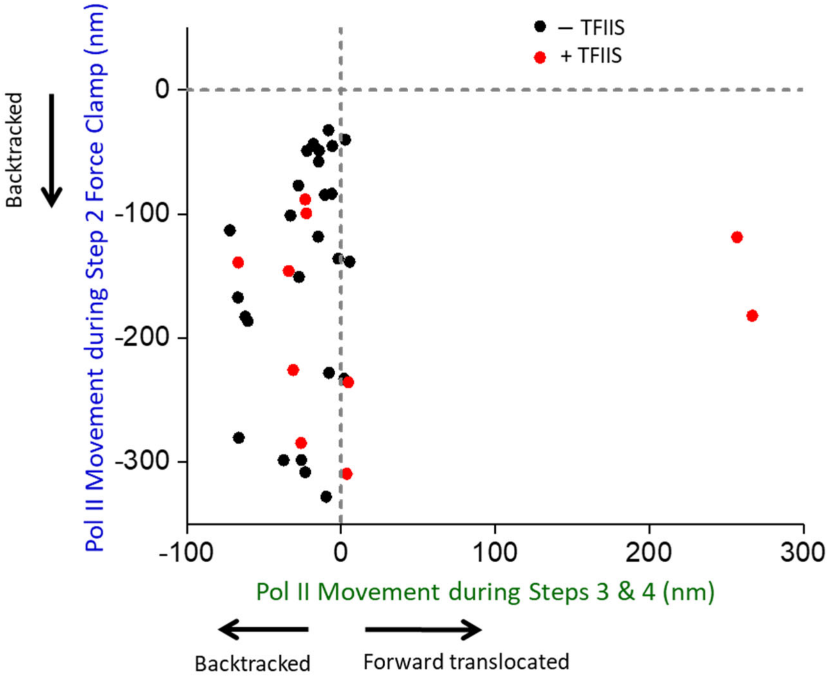


**Supplementary Figure 5.** (+)/(-) TFIIS transcription after hybrid formation

Measured data for all traces that formed an RNA-DNA hybrid after Pol II backtracking. The backtrack during the force clamp is plotted against Pol II movement in steps 3 (rezipping) and 4 (unzipping). (-) TFIIS data are shown in black (*N* = 26) and (+) TFIIS data are shown in red (*N* = 10). After extensive backtracking, Pol II should be unable to resume transcription unless TFIIS is present to encourage RNA cleavage. 2 of 10 (+) TFIIS traces exhibit Pol II elongation after extensive backtracking.

**Supplementary Table 1.** Primer and oligo sequences.

| Oligo Name | Sequence (5' → 3') |
|---|---|
| RNA 14 | UUUUUAUCGAGAGG (Underlined sequence complementary to the gap) |
| ARP 77 | /5Phos/TCAGCCCTATAGGATACTTACAGCCATCGAGAGGGACAAG GCGAATACCCATCCCAATCGGCCTGCTGGTGACACC |
| US NTS | /5Phos/GTCCTAATTCGAGCTCGGTACCCGGGGATCC |
| DS NTS | /5Phos/TCAGCGGATCCTCTAGAGTCCTTCAGCGAT |
| TS | CTGAAGGACTCTAGAGGATCCGCTGAGGTGTCACCAGCAGGCCGA TT |
| 0.65/1.4 kb CD/HO upstream PCR F | GCTCCTGTCGTTGAGGACCC |
| 0.65 kb CD upstream PCR R | GCTTACAGACACCTAGTGACCG |
| 1.4 kb HO upstream PCR R | CTGCTAATCCTGTTACCAGCTACTGC |
| 1.5 HO/CD Downstream PCR F | AATACTGTTCACCAACGATCTGGATCACG |
| 1.5 HO/CD Downstream PCR R | GGGACACACACGCCAGCTACTG |
| Daughter Strand PCR F | CGCGTTTCGGTGATGACGGTGA |
| Leading Strand PCR R_Bio | /5BiosG/TACCGATGAAACGAGAGAGGATGC |
| Lagging Strand PCR R_Dig | /5DiGN/TACCGATGAAACGAGAGAGGATGC |
| Upper Leading Strand | /5Phos/GGGACAGACGCTGTCCGCGCCAGTGCAGAATAAGGAGTC ATTCGTGGGGTGGAC |
| Upper Leading Strand_invdT | /5Phos/GGGACAGACGCTGTCCGCGCCAGTGCAGAATAAGGAGTC ATTCGTGGGG/3invdT/ |
| Lower Leading Strand | /5Phos/ACCCAGATGCGTGTGCGTAGAGCGGACCGCGTCCACCCCA CGAATGACTCCTTATTCTGCACTGGCGCGGACAGCGTCTG |
| Upper Lagging Strand | CAGCGCCAGACTGGGGGCGTCCTGCAGAAGGCTCCCACGACGACA CCGAC |
| Lower Lagging Strand | /5Phos/GGGAGTCGGTGTCGTCGTGGGAGCCTTCTGCAGGACGCC CCCAGTCTGGCGCTGGCGGTCCGCTCTACGCACACGCATCTGGGTC TA |

#### References


1 Komissarova, N., Kireeva, M. L., Becker, J., Sidorenkov, I. & Kashlev, M. in *Methods in Enzymology* Vol. 371 233-251 (Academic Press, 2003).

2 Christie, K. R., Awrey, D. E., Edwards, A. M. & Kane, C. M. Purified yeast RNA polymerase II reads through intrinsic blocks to elongation in response to the yeast TFIIS analogue, P37. *J Biol Chem* **269**, 936-943 (1994).

3 Christie, K. R., Awrey, D. E., Edwards, A. M. & Kane, C. M. Purified Yeast Rna Polymerase-Ii Reads through Intrinsic Blocks to Elongation in Response to the Yeast Tfiis Analog, P37. *J Biol Chem* **269**, 936-943 (1994).

4 Inman, J. T. *et al.* DNA y structure: a versatile, multidimensional single molecule assay. *Nano Lett* **14**, 6475-6480 (2014).

5 Sun, B. & Wang, M. D. Single-Molecule Optical-Trapping Techniques to Study Molecular Mechanisms of a Replisome. *Methods Enzymol* **582**, 55-84 (2017).

6 Killian, J. L., Inman, J. T. & Wang, M. D. High-Performance Image-Based Measurements of Biological Forces and Interactions in a Dual Optical Trap. *ACS Nano* **12**, 11963-11974 (2018).

7 Sun, B. *et al.* Helicase promotes replication re-initiation from an RNA transcript. *Nat Commun* **9**, 2306 (2018).

8 Hall, P. M. *et al.* Polarity of the CRISPR roadblock to transcription. *Nat Struct Mol Biol* **29**, 1217-1227 (2022).

9 Ye, F., Inman, J. T., Hong, Y., Hall, P. M. & Wang, M. D. Resonator nanophotonic standing-wave array trap for single-molecule manipulation and measurement. *Nat Commun* **13**, 77 (2022).

10 Le, T. T. *et al.* Etoposide promotes DNA loop trapping and barrier formation by topoisomerase II. *Nat Chem Biol* **19**, 641-650 (2023).

11 Walter, W., Kireeva, M. L., Studitsky, V. M. & Kashlev, M. Bacterial Polymerase and Yeast Polymerase II Use Similar Mechanisms for Transcription through Nucleosomes*. *Journal of Biological Chemistry* **278**, 36148-36156 (2003).

12 Kireeva, M. L. *et al.* Nature of the nucleosomal barrier to RNA polymerase II. *Mol Cell* **18**, 97-108 (2005).

13 Kireeva, M. L. *et al.* Nucleosome remodeling induced by RNA polymerase II: loss of the H2A/H2B dimer during transcription. *Mol Cell* **9**, 541-552 (2002).

14 Le, T. T. *et al.* Mfd Dynamically Regulates Transcription via a Release and Catch-Up Mechanism. *Cell* **172**, 344-357.e315 (2018).

15 Johnson, D. S., Bai, L., Smith, B. Y., Patel, S. S. & Wang, M. D. Single-molecule studies reveal dynamics of DNA unwinding by the ring-shaped T7 helicase. *Cell* **129**, 1299-1309 (2007).

16 Wang, M. D., Yin, H., Landick, R., Gelles, J. & Block, S. M. Stretching DNA with optical tweezers. *Biophysical Journal* **72**, 1335-1346 (1997).
